# cAb-Rep: A Database of Curated Antibody Repertoires for Exploring antibody diversity and Predicting Antibody Prevalence

**DOI:** 10.1101/765099

**Authors:** Yicheng Guo, Kevin Chen, Peter D. Kwong, Lawrence Shapiro, Zizhang Sheng

## Abstract

The diversity of B cell receptors provides a basis for recognizing numerous pathogens. Antibody repertoire sequencing has revealed relationships between B cell receptor sequences, their diversity, and their function in infection, vaccination, and disease. However, many repertoire datasets have been deposited without annotation or quality control, limiting their utility. To accelerate investigations of B cell immunoglobulin sequence repertoires and to facilitate development of algorithms for their analysis, we constructed a comprehensive public database of curated human B cell immunoglobulin sequence repertoires, cAb-Rep (https://cab-rep.c2b2.columbia.edu), which currently includes 306 immunoglobulin repertoires from 121 human donors, who were healthy, vaccinated, or had autoimmune disease. The database contains a total of 267.9 million V(D)J heavy chain and 72.9 million VJ light chain transcripts. These transcripts are full-length or near full-length, have been annotated with gene origin, antibody isotype, somatic hypermutations, and other biological characteristics, and are stored in FASTA format to facilitate their direct use by most current repertoire-analysis programs. We describe a website to search cAb-Rep for similar antibodies along with methods for analysis of the prevalence of antibodies with specific genetic signatures, for estimation of reproducibility of somatic hypermutation patterns of interest, and for delineating frequencies of somatically introduced *N*-glycosylation. cAb-Rep should be useful for investigating attributes of B cell sequence repertoires, for understanding characteristics of affinity maturation, and for identifying potential barriers to the elicitation of effective neutralizing antibodies in infection or by vaccination.

## Introduction

B cells comprise a crucial component of the adaptive immune response (Murphy, 2014). B cells recognize three-dimensional epitopes of antigens through the variable domains of the B cell receptor (BCR), or its various secreted forms of antibody. The variable domains of BCRs and antibodies are composed of immunoglobulin heavy and light chains, encoded by separate genes. Through V(D)J gene recombination and somatic hypermutation (SHM), a high level of sequence diversity is introduced to the variable domain (Briney et al., 2019; Elhanati et al., 2015; Murphy, 2014; Soto et al., 2019), allowing B cells to recognize diverse antigens. Thus, an interrogation of B cell diversity and function is key to understanding the B cell immune response. The application of next-generation sequencing (NGS) to BCR repertoires provides snapshots of BCR diversity, and such studies in the past decade have characterized numerous features of B cell responses to infection, immunization, and autoimmune disease (Elhanati et al., 2015; Francica et al., 2015; Galson et al., 2016a; Galson et al., 2015; Galson et al., 2016b; Joyce et al., 2016; Kwong et al., 2017; Nielsen and Boyd, 2018; Rubelt et al., 2016; Sheng et al., 2016; Sheng et al., 2017; Stern et al., 2014; Tipton et al., 2015; Vander Heiden et al., 2017; Wu et al., 2015). Databases, programs, and websites have been developed to store, process, and annotate BCR repertoire data (Brochet et al., 2008; DeWitt et al., 2016; Margreitter et al., 2018; Miho et al., 2018; Schramm et al., 2016; Vander Heiden et al., 2018; Vander Heiden et al., 2017) (software reviewed in (Chaudhary and Wesemann, 2018)). Nonetheless, most repertoire data are deposited in public databases in the format of raw NGS reads, which contain both sequencing duplicates and sequencing errors and may be difficult to annotate. A comprehensive database of curated and well-annotated BCR transcripts, should therefore accelerate repertoire studies including but not limited to characterization of B cell receptor diversity, mechanisms of clonal expansion, development of BCR repertoire analysis algorithms, estimation of prevalence of antigen-specific antibodies and their precursors, and functions of SHM. In addition, such a database should assist researchers in performing repertoire-related data mining.

Designing vaccines that can elicit broadly neutralizing antibodies (bnAbs) is a long-term goal for preventing infections from fast evolving and/or highly diversified pathogens such as HIV-1 and influenza (Burton and Mascola, 2015; Corti and Lanzavecchia, 2013). Studies have revealed that some epitopes could elicit bnAbs in different individuals with similar modes of recognition and shared genetic signatures (e.g. V(D)J gene origin, complementarity determining region 3 (CDR3) length, CDR3 motifs, SHM) (Andrabi et al., 2015; Ekiert et al., 2009; Joyce et al., 2016; Kwong and Mascola, 2018; Wu et al., 2011; Zhou et al., 2015; Zhou et al., 2013), defined as convergent, multidonor or public bnAb classes. Such antibody classes often target conserved epitopes; those that can activate and mature precursor B cells with bnAb potential might provide templates for universal vaccines (Chuang et al., 2019; Kwong and Mascola, 2018; Kwong and Wilson, 2009; Pappas et al., 2014). However, not all antibody classes appear with high frequencies in BCR repertoires; those that do not may be limited due to disfavored developmental steps or “roadblocks”. Thus, the identifications of bnAb classes with precursor-like cells prevalent in humans is a critical consideration for determining which template antibodies are good targets for elicitation, and thus for immunogen design (Dosenovic et al., 2018; Jardine et al., 2016a; Joyce et al., 2016). A comprehensive database of curated B cell repertoire transcripts, for which uncertainties and variations in B cell repertoires (e.g. sampling bias, infection history, aging, and genetic diversification)(Rubelt et al., 2016) have been minimized, would be helpful for predicting antibody class prevalences.

A database of curated BCR repertoires could, moreover, be used to investigate SHM preference and mechanisms of affinity maturation. Antibodies accumulate mutations with high preference determined by both intrinsic gene mutability and functional selection (Murphy, 2014; Odegard and Schatz, 2006). We previously developed gene-specific substitution profiles (GSSPs) to characterize positional substitution types and frequencies in 69 human V genes (Sheng et al., 2017). By incorporating additional B cell transcripts, GSSPs could be built for more genes, and this could have broad application; GSSPs or similar approaches have been applied to examine whether bnAbs mature with shared pathways (Bonsignori et al., 2016), to identify highly frequent SHMs and common mechanisms of function (Koenig et al., 2017; van de Bovenkamp et al., 2018), and to estimate whether rare SHMs (SHMs generated with very low frequency by the SHM machinery) in bnAbs could form barriers to re-elicitation by vaccination (Chuang et al., 2019).

In this study, we constructed a database of curated human B cell immunoglobulin sequence repertoires (cAb-Rep) from 306 high quality human repertoires, and developed methods to search cAb-Rep using either sequence or sequence signature. We used the database to construct GSSPs for 102 human antibody V genes and developed a script to identify rare SHMs in input sequences. We evaluated the robustness of query results relative to the number of repertoires, using antibody prevalence estimates as a test case. In summary, the database developed here should help to investigate B cell repertoire features, to find antibody templates for vaccine design, and overall to understand mechanisms of antibody development.

## Materials and Methods

### BCR repertoire dataset

We assembled a total of 376 BCR repertoire datasets from 108 human donors deposited in the NCBI short reads archive (SRA) database (https://www.ncbi.nlm.nih.gov/sra). These repertoires were all sequenced using illumina MiSeq or HiSeq and were published in 9 studies (Table 1). In addition, annotated full length sequences of three donors (sequenced by AbHelix with high depth or HD repertoires, ∼10^8^ high quality sequences per donor) (Soto et al., 2019) and the near full length V(D)J sequences (sequenced with framework 1 region primers) from ten HD repertoires (Briney et al., 2019) were downloaded from the links provided by the authors.

**Table 1:**
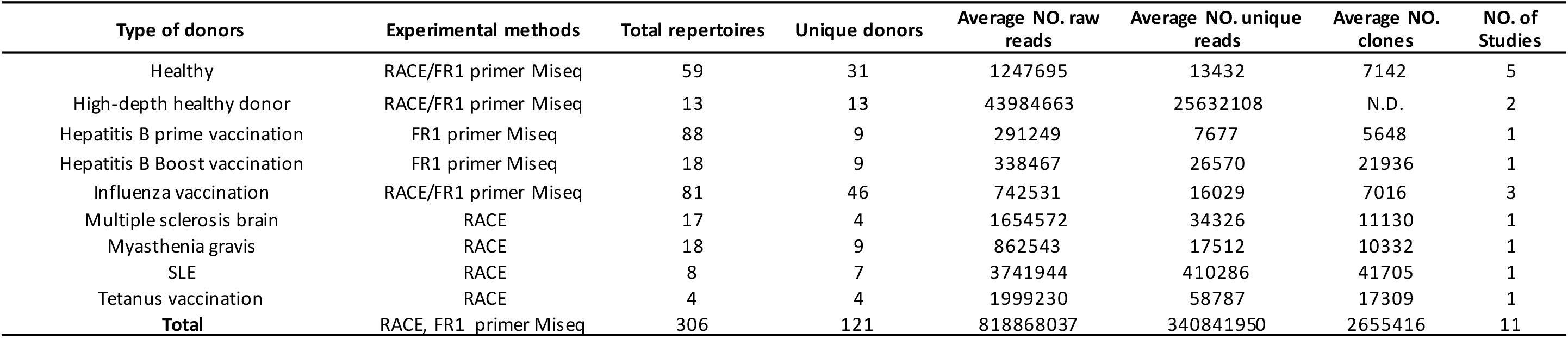
Summary of datasets, type of donors, experimental methods, unique reads, raw reads, number of clones, and references

### Next-generation sequencing data processing

The NGS data were analyzed with the SONAR pipeline, version 2.0 (https://github.com/scharch/sonar/) developed in our lab (Schramm et al., 2016). Briefly, USEARCH was used to merge the 2×300 raw reads to single transcripts and to remove transcripts potentially containing more than 20 miscalls calculated from sequencing quality score (Edgar and Flyvbjerg, 2015). Merged transcripts shorter than 300 nucleotides were removed. BLAST (http://www.ncbi.nlm.nih.gov/blast/) was used to assign germline V, D, and J genes to each transcript with customized parameters (Altschul et al., 1997; Schramm et al., 2016). CDR3 was identified from BLAST alignment using the conserved 2^nd^ Cysteine in V region and WGXG (heavy chain) or FGXG (light chain) motifs in J region (X represents any of the 20 amino acids). To assign an isotype for each heavy chain transcript, we used BLAST with default parameters to search the 3’ terminus of each transcript against a database of human heavy chain constant domain 1 region obtained from the international ImMunoGeneTics information system (IMGT) database. A BLAST E-value threshold of 1E-6 was used to find significant isotype assignments. Then, sequences other than the V(D)J region of a transcript were removed and transcripts containing frame-shift and/or stop codon were excluded.

We then applied two approaches to remove PCR duplicates and sequencing errors. For datasets sequenced with unique molecular identifier (UMI), we first grouped merged raw transcripts having identical UMI. Due to PCR crossover and UMI collisions, transcripts in a group may originate from different cells (Briney et al., 2019; Rubelt et al., 2016). We therefore clustered the raw transcripts in each group using USEARCH with a 97% sequence identity. All transcripts in a USEARCH cluster were aligned using CLUSTALO (Sievers et al., 2011) and a consensus sequence was generated. To further reduce redundant transcripts derived from multiple mRNA molecules of the same cell, we removed duplicates in the consensus sequences. Singletons, which have neither UMI duplicates nor consensus duplicates, were removed. SONAR was used to annotate the unique transcripts. For datasets don’t contain UMI, similar to our previous studies (Schramm et al., 2016; Sheng et al., 2017; Wu et al., 2015), we clustered transcripts of each repertoire using USEARCH with sequence identity of 0.99, and only one transcript with the highest sequencing depth or numbers of duplicates was selected from each cluster for later analyses. To further remove low quality reads, we excluded clusters with size less than 2. Finally, a unique dataset of transcripts was generated for each repertoire. To reduce artifacts and effect of small sample size on the comparisons of features of antibody repertoires, we excluded repertoires containing fewer than 1700 unique transcripts.

For each of the 13 HD repertoires curated in previous studies, we further removed duplicates, sequences containing frameshifts or/and stop codons, and corrected annotation errors to form a unique dataset. The gene annotation information was retrieved from the annotation files provided by the studies.

### Germline gene database and prediction of new gene/alleles

The human germline gene database from IMGT was used to assign germline V(D)J genes. To detect new germline genes or alleles, we combined unique reads from 108 repertoires, and used IgDiscover v0.9 with default parameters to identify potential new germline genes or alleles that are observed in multiple repertoires (Corcoran et al., 2016). While IgDiscover prefers to identify new genes or alleles from IgM repertoire, we used repertoires containing all isotypes. Nonetheless, identical unique reads were observed for each predicted gene/allele in at least two donors, suggesting that the predictions are still reliable. The predicted genes were submitted to European Nucleotide Archive (ENA) with project accession numbers: PRJEB31020. For each of the 13 HD repertoires, we randomly selected 1 million IgM transcripts for novel gene prediction (as recommended by IgDiscover manual) and no new gene or allele was found.

### V and J gene usage and antibody position numbering

The unique dataset for each repertoire was used to calculate the distributions of germline gene usage and SHM levels. For each transcript, we used ANARCI to assign each position according to the Kabat, Chothia, and IMGT numbering schemes (Dunbar and Deane, 2016).

### Signature prevalence and rarefaction analysis

We developed a python script, Ab_search.py, to search the BCR repertoire database in two modes: signature motif and full V(D)J sequence. Briefly, in the signature searching mode, an amino acid signature motif from either CDR3 region or other positions of interest (defined with the Kabat, Chothia, and IMGT numbering scheme) is converted to python regular expression format and searched against all transcripts in the database. For the sequence searching mode, BLAST+ is called to find similar transcripts with E-value less than 1E-6 (Camacho et al., 2009). Signature frequency in each repertoire is calculated by dividing the number of matched transcripts with the total number of unique transcripts originated from the same germline gene or allele.

To evaluate the effect of number of repertoires on estimation of signature frequency, we performed rarefaction analysis by sampling *i* subset repertoires from PBMC samples (1 to 35). For each repertoire size *i*, the random sampling was performed N times (we used N=20 in the current study) and the mean signature frequency from each sampling 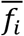 was calculated. Then the coefficient of variance for each *i*, *CV_i_*, was calculated as:

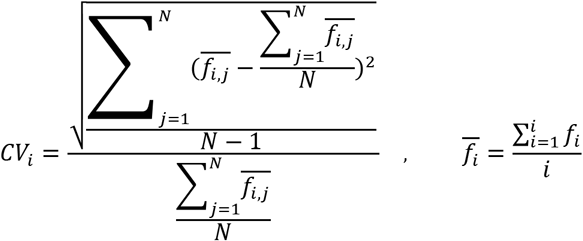

#### Identification of antibody clones and Construction of GSSPs

For each repertoire, we first sorted all transcripts to groups based on identical V and J gene usages. For each group, transcripts with 90% CDR3 sequence identity and the same CDR3 length were clustered into clones using USEARCH. One representative sequence was selected in each clone and representative transcripts from all repertoires were combined to build GSSPs for V genes using mGSSP (Sheng et al., 2017). We exclude GSSPs built with less than 100 clones because of lack of information (Sheng et al., 2017). A python script, SHM_freq.py, was developed to identify SHMs in an input sequence, to search the GSSP of an assigned V gene, and to output the rarity of each mutation, calculated as: Rarity = (1 – Frequency of mutation) * 100%.

#### Construction of gene-specific N-glycosylation profile

Glycosylation sites for sequences having SHMs greater than 1% in the unique datasets of healthy and vaccination donors were predicted using an artificial neural network method NetNGlyc v1.0 (http://www.cbs.dtu.dk/services/NetNGlyc/) (R. Gupta et al., 2004). Predictions with high specificity (potential greater than 0.5 and jury agreement of 9/9) and do not match Asn-Pro-Ser/Thr motifs were included. N-glycosylation sites encoded in germline genes were excluded.

#### Statistical tests

ANOVA and T tests were performed in R.

### Results

#### cAb-Rep contains a diverse compendium of annotated next-generation sequencing datasets

We first assembled 376 BCR repertoire deep sequencing data sets from NCBI SRA database, each sequenced by Illumina MiSeq or HiSeq and with library preparation protocols to cover full length V(D)J region (5’ primers at leader regions or 5’ Rapid amplification of cDNA ends (RACE), 3’ primers targeting constant region 1) or near full length (5’ primers at N-terminus of framework 1 region). These datasets contain a total of 247 million raw reads (Table 1 and S1). We performed quality control including removing sequencing errors, PCR crossover, PCR duplicates, and annotated each sequence using SONAR (See Methods and Fig.1). A total of 293 repertoires from 108 donors were selected to construct the cAb-Rep database, with 6.6 and 0.9 million curated full length V(D)J heavy and light chain transcripts. 25541 curated transcripts were obtained per repertoire. In addition, annotated sequences from 13 healthy donor repertoires sequenced with high depth (more than 20 million unique sequences from about 10^7^ to 10^8^ B cells per donor) (Briney et al., 2019; Soto et al., 2019), were curated and incorporated.

**Figure 1.**
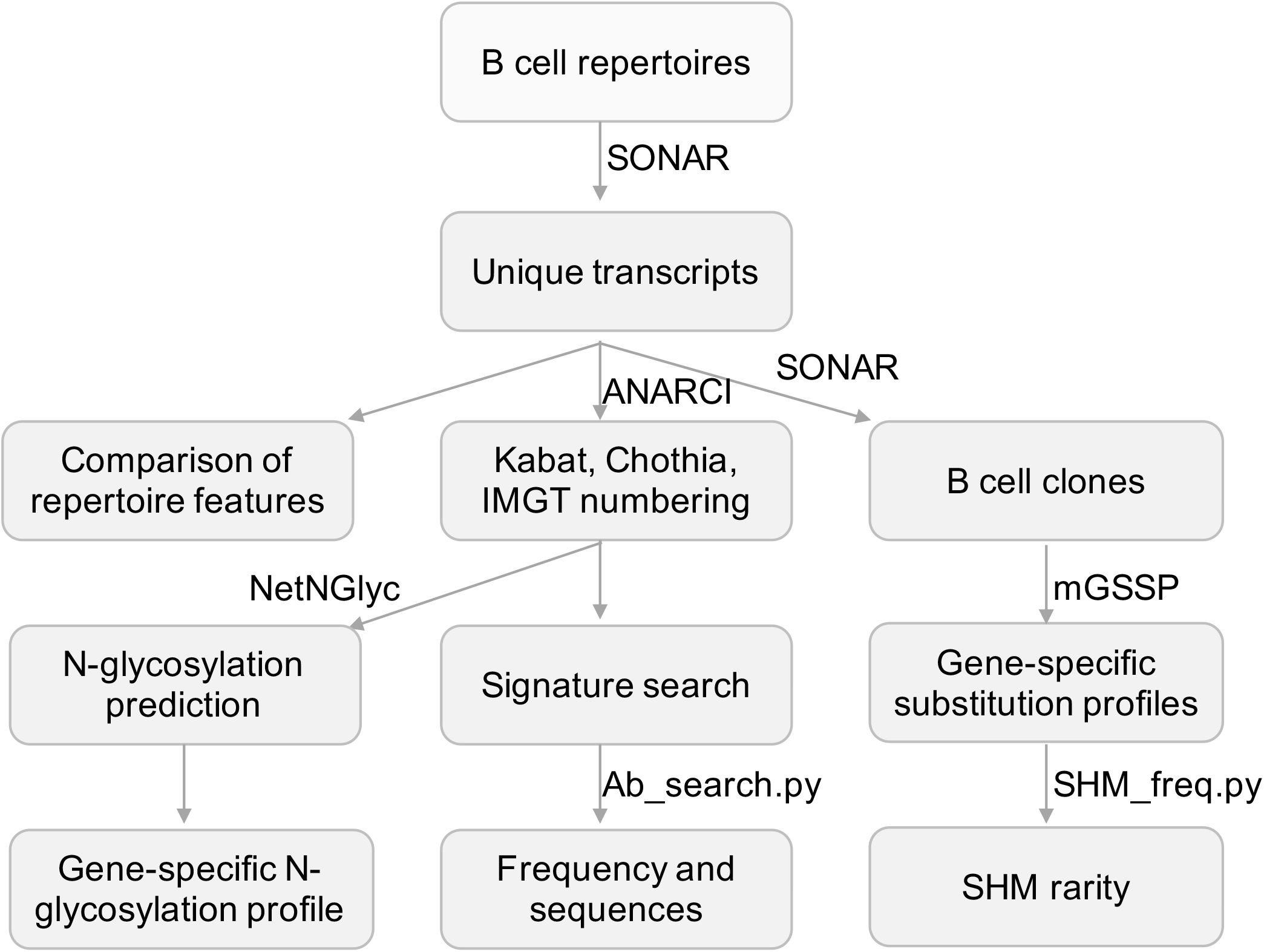
Flowchart for the processing of repertoire data and developed tools. Next-generation sequencing data was processed and annotated using SONAR. Other published programs used were highlighted in bold font. Scripts developed in this study were in italic font.

These repertoires are from studies including Hepatitis B vaccination, influenza vaccination, Tetanus vaccination, Multiple Sclerosis, Myasthenia Gravis, Systemic Lupus Erythematosus, and healthy donors (Table 1 and S1) (Galson et al., 2016a; Galson et al., 2015; Galson et al., 2016b; Gupta et al., 2017; Joyce et al., 2016; Rubelt et al., 2016; Stern et al., 2014; Tipton et al., 2015; Vander Heiden et al., 2017). Among all repertoires, 72 repertoires are from 43 healthy donors (35.5% of total donors). The database repertoiores were obtained from three tissues: 2 brain samples, 15 cervical lymph node samples, and 289 peripheral blood mononuclear cell (PBMC) samples. Among PBMC samples, 16 were from antibody secreting B cells, 27 from Hepatitis B surface antigen (HBsAg)+ B cells, 8 from Human Leukocyte Antigen – DR isotype (HLA-DR)^+^ plasma cells, 139 from IgG^+^ B cells, 20 from memory B cells, 34 from IgM or naïve B cells, and 52 from whole PBMC (Table S1). Thus, cAb-Rep contains BCR repertoire data in different settings, which will enable cross-study comparisons.

For each transcript in cAb-Rep, we numbered positions using Kabat, Chothia, and IMGT schemes (Fig. 1), and annotated gene origin, CDR3 sequence and length, somatic hypermutation level, isotype, donor, and clonotype. These curated datasets can be used for downstream analyses. The transcripts are stored in FASTA format with annotation information in the header line of each transcript. Such a format can be easily altered to feed into other programs for repertoire analysis.

Because the 13 HD repertoires contains over 330 million unique sequences, to avoid sampling bias for profile constructions and improve speed of searching cAb-Rep (see sections below), we randomly selected 20,000 unique sequences of each isotype of each repertoire and combined with the rest of repertoires for all analyses below. Nonetheless, in case rare events of BCR signatures are of interests, we provided an option in our scripts and at cAb-Rep website to search the 13 HD repertoires alone.

#### Ab_search identifies similar antibody transcripts and estimates signature frequency

To identify antibody transcripts of interest from cAb-Rep, we developed a python script, Ab_search.py. The script accepts sequences or amino acid motifs as input and output frequency of the signature and matched sequences (See Materials and Methods). Here, as examples, we searched transcripts having signature motifs of a selected malaria antibody class in PBMC BCR repertoires or of a selected HIV-1 bnAb classes in IgM and IgG repertoires (Fig.2).

**Figure 2.**
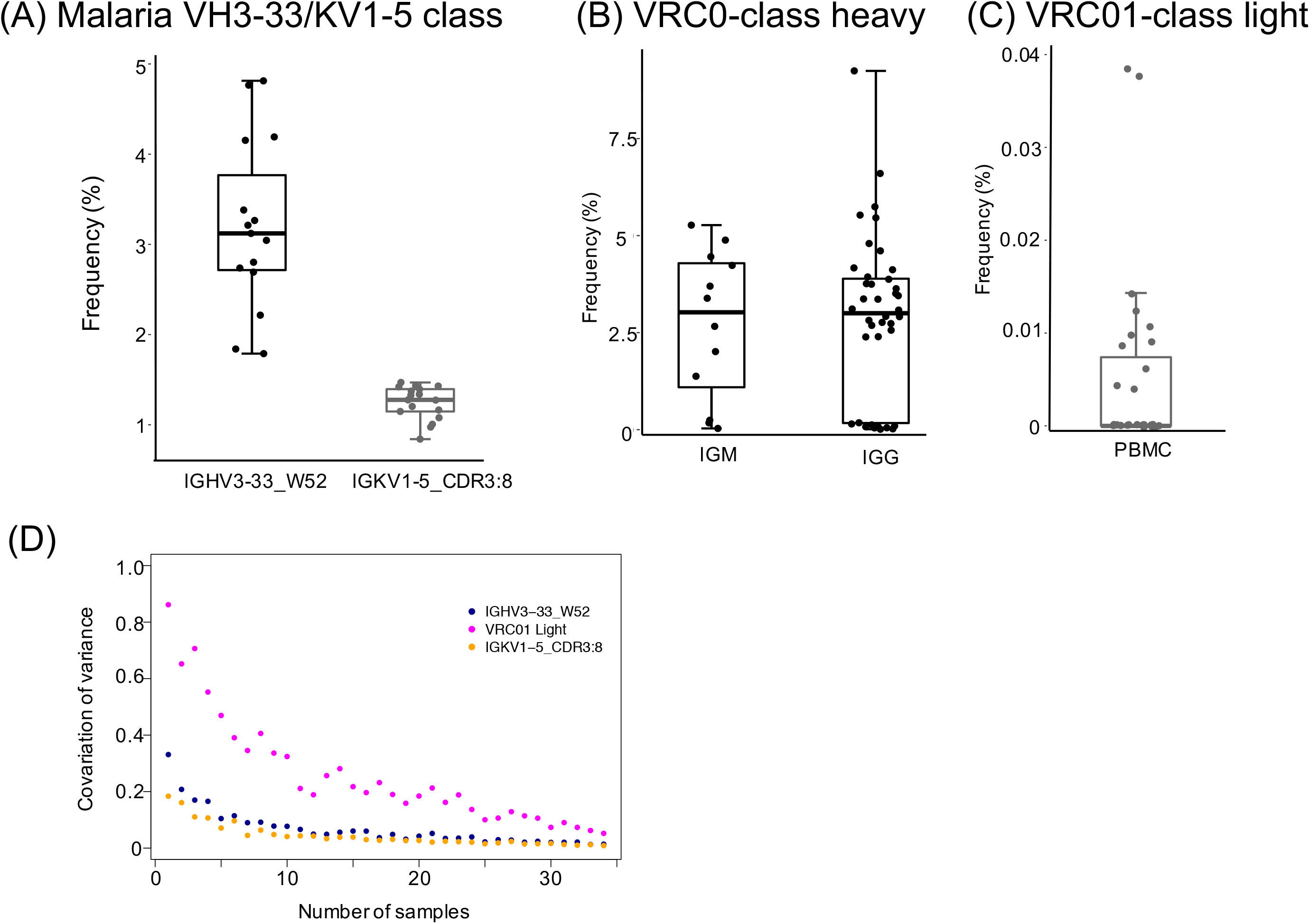
Frequencies of malaria and HIV-1 broadly neutralizing antibody classes-like transcripts. (A) Frequencies of influenza IGHV3-33/IGKV1-5 class heavy and light chain signatures in 15 heavy and 17 light chain PBMC repertoires of healthy donors. Signatures are: IGHV3-33 gene and W52 for heavy chain and IGKV1-5 and 8 amino acids CDRL3 for light chain. (B) VRC01 class heavy chain and light chain signatures were showed in (B) and (C). The searched signatures were IGHV1-2*02 origin and either IGKV1-33, IGKV3-15, IGKV3-20, or IGLV2-14 origin plus a CDR3 of five amino acids matching motif X-X-[AFILMYWV]-[EQ]-X for heavy and light chains respectively. (D) Rarefaction analysis of the malaria heavy and VRC01 light signatures from 35 PBMC repertoires showed the variations of signature frequencies between random samplings (measured with coefficient of variation) reduce when more than 10 repertoires were sampled.

The selected malaria antibody class contains both heavy (IGHV3-33 gene with a tryptophan at position 52) and light chain (IGKV1-5 gene with 8 amino acid CDRL3) signatures (Imkeller et al., 2018). The heavy chain signature was observed with a frequency of 3.2% ± 0.94% in 15 PBMC repertoires in healthy donors (Fig.2A), the light chain signature was found with a frequency of 1.24% ± 0.18% in 17 healthy donor PBMC repertoires. All healthy donors we searched contain both heavy and light sequences. By assuming random pairing (Jayaram et al., 2012), heavy-light paired antibodies similar to the malaria antibody class could be generated with a high frequency (approximately 40 per million B cells). Thus, vaccine mediated elicitation of such antibodies are promising candidates to enable protection.

The HIV-1 VRC01 class contains both heavy and light chain signatures (West et al., 2012; Wu et al., 2011; Zhou et al., 2015; Zhou et al., 2013). Its heavy chain uses IGHV1-2*02 allele. The light chain is originated from a limited set of genes, including IGKV1-33, IGKV3-15, IGKV3-20, or IGLV2-14 and contains a CDR3 of five amino acids matching motif X-X-[AFILMYWV]-[EQ]-X. Searching cAb-Rep with these signatures showed that the heavy chain allele for VRC01 appears in IgM and IgG repertories with similar frequencies (2.7%±1.9% and 2.85%±2.13% respectively) (Fig. 2B). Further, VRC01 light chain-like transcripts were found in ∼33% of donors, with overall frequencies of 0.005%±0.0098% in PBMCs (Fig. 2C). By assuming random pairing of heavy and light chains, our calculation of the mean frequency of VRC01 class-like antibodies was 1.4 per million B cells in healthy donors, close to estimates from previous studies (Jardine et al., 2016a; Zhou et al., 2013).

#### Impact of repertoire diversity on signature prevalence

Due to diversification in BCR repertoires by antigen selection and other factors, the predicted prevalence of transcripts of interest could vary. To evaluate the effect of the number of sampled repertoires on prevalence prediction, we performed rarefaction analysis to sample a subset of repertoires (ranging from 1 to 35) with each subset randomly sampling 20 times to calculate frequencies of the malaria IGHV3-33/IGKV1-5 class and VRC01 class light chain signatures. For each sampling size, we calculated the coefficient of variation (standard deviation/mean) to measure the degree of variation among sampling repeats (Fig. 2D). For both signatures, our analysis revealed that the coefficient of variation decreases dramatically when the sampling size increases to 10 or more. This suggests that 10 or more repertoires will be optimal to have a consistent estimation of signature frequency.

#### Gene-specific substitution profiles and substitution frequency analysis

To investigate substitution preferences in V genes, we predicted new germline genes in the database, clustered transcripts in each repertoire into clones, and selected one representative sequence per clone to build gene-specific substitution profiles (GSSPs) (see Methods). Overall, we identified 5 novel heavy chain alleles, 2 novel lambda chain alleles, and 1 kappa chain allele, each of which was found in two or more donors. By incorporating germline gene sequences in database and prediction, we built GSSPs for 102 human V genes, compared to GSSPs built for 69 genes using three repertoires in our previous study. Our analysis further showed that the GSSPs of the two studies are highly consistent (r = 0.983 for IGHV1-2 gene, Fig. 3A).

**Figure 3.**
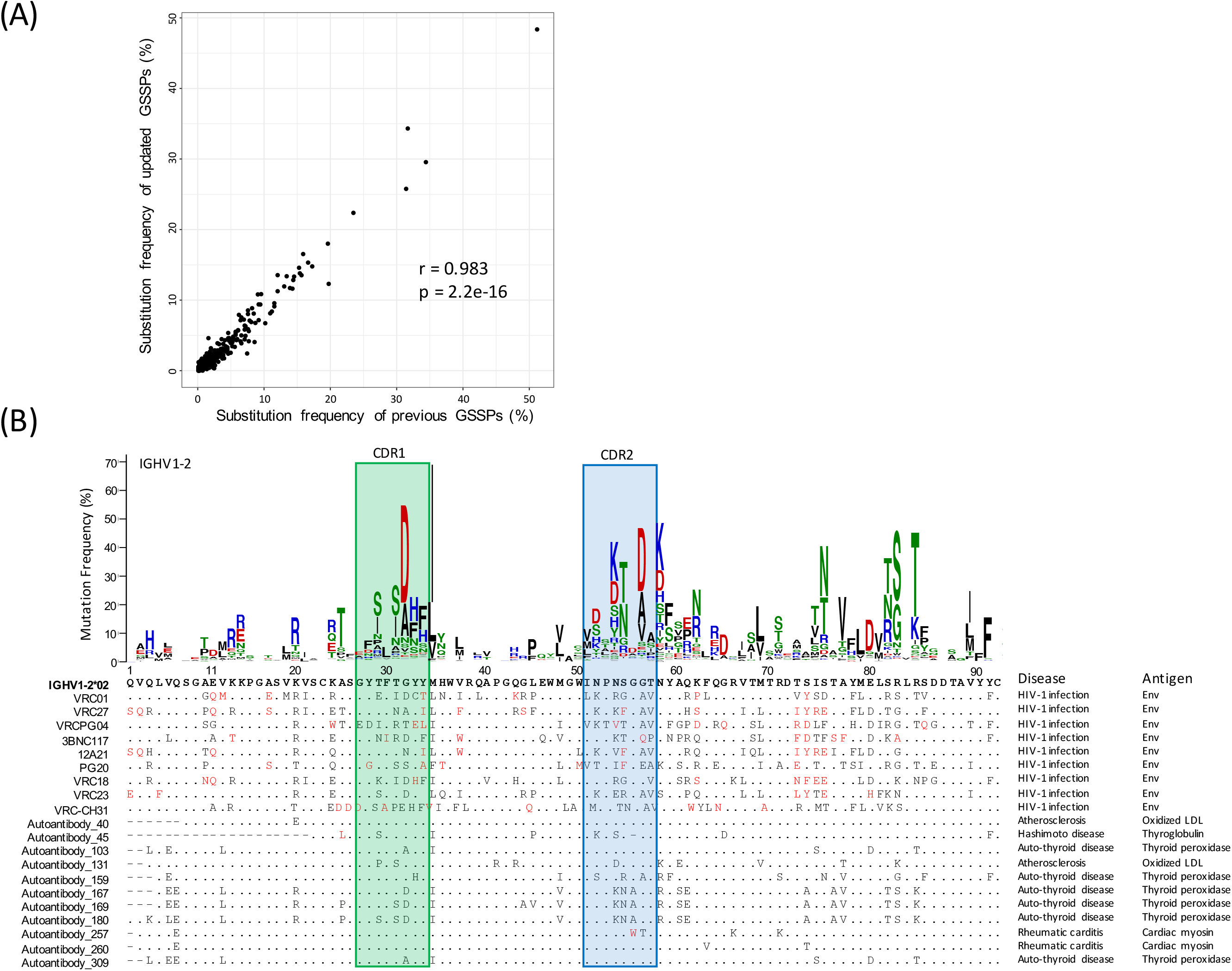
Comparison of gene-specific substitution profiles and usage of a substitution profile for investigating substitution preference. (A) Comparison of substitution frequencies of all amino acid types at all IGHV1-2 positions estimated using cAb-Rep dataset and previous dataset. A Pearson correlation coefficient of 0.982 suggested that the substitution profiles of IGHV1-2 are highly consistent. (B) The gene-specific substitution profile of IGHV1-2 and rarity of somatic hypermutations in HIV-1 bnAbs and autoantibodies. Rare mutations, colored red, are observed frequently in HIV-1 bnAbs but not in autoantibodies, suggesting the mutation patterns in HIV-1 bnAbs may be generated with low frequency. For each antibody sequence, residues identical to IGHV1-2*02 germline gene were shown with dots. Missing residues were showed with minus sign. The disease and antigen were labeled on the right side of each sequence.

To facilitate exploring substitution preference, we developed a python script, SHM_freq.py, to identify mutations in an input sequence, call the GSSP of corresponding V gene, and find the frequency of the mutation being generated by the somatic hypermutation machinery. To demonstrate how this information can be helpful, we analyzed frequencies of substitutions observed in the heavy chain of VRC01 class bnAbs (Fig. 3B). This analysis showed that all lineages in this class contain over 30% mutations, with ∼30% of the mutations being low frequency or rare mutations (frequency <0.5% in IGHV1-2 GSSP). Functional studies have shown that some rare mutations are essential for function (Jardine et al., 2016b). However, since rare mutations are generated with low frequency, the likelihood of immunogens maturing antibodies to have similar mutations could be low or require longer maturation times. In contrast, we observed that autoantibodies (e.g. collected from HIV, autoimmune thyroid disease, atherosclerosis, Hashimoto disease, and rheumatic carditis (Chazenbalk et al., 1993; Hexham et al., 1992; Jeon et al., 2007; Pichurin et al., 2001; Pichurin et al., 2002; Wu et al., 1998)) originated from IGHV1-2 genes contain very few rare mutations, suggesting somatic mutations may not provide a barrier to elicitation of these lineages.

#### Gene-specific N-glycosylation profiles (GSNPs)

Post-translation modifications (PTM) (glycosylation, tyrosine sulfation, etc.), which affects antibody functions (Doria-Rose et al., 2014; van de Bovenkamp et al., 2018), can be introduced to antibodies by V(D)J recombination and somatic hypermutation processes. To understand the frequency and preference of PTMs generated by somatic hypermutation, as an example, we predicted V-gene-specific frequency of N-glycosylation sequons at each position using healthy and vaccination donor unique sequences that having more than 1% SHM. Overall, consistent with previous study (van de Bovenkamp et al., 2018), the predicted N-glycosylation sites are enriched in CDR1, CDR2, and framework3 regions, but different genes have different hotspots for glycosylation (Fig. 4A). Structural analysis showed that the side chains of these hotspot positions are surface-exposed (Fig. 4B), suggesting these sites are spatially accessible for modification. We think the GSNPs provide information for further experimental validation and investigations of functions of N-glycosylations.

**Figure 4.**
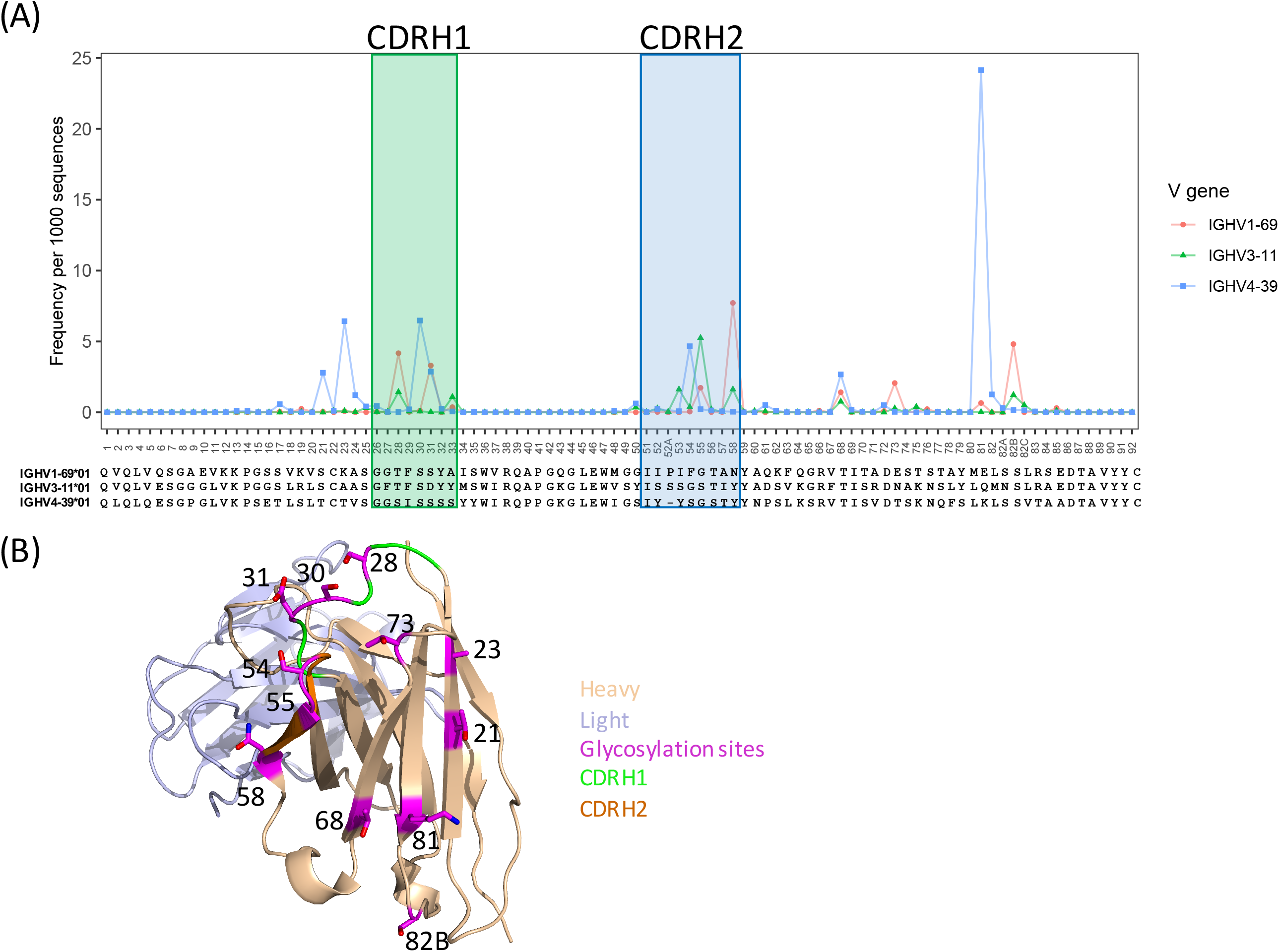
Predicted glycosylation sites in V genes and structural location. (A) Hotspots of glycosylation sites in IgHV1-69, IgHV3-11, and IgHV4-39 genes. (B) A structural demo (PDBID: 1dn0) to show that the predicted glycosylation hotspots are surface-exposed, suggesting they are accessible for post-translational modification.

#### cAb-Rep website to search frequencies of signature motif and SHM

While we developed scripts to search cAb-Rep, these may not be friendly to users not familiar with programing. Therefore, we developed a website for searching cAb-Rep (https://cab-rep.c2b2.columbia.edu/). The website implements all functions of the scripts we developed above, including querying cAb-Rep using the three signature modes (CDR3, position, BLAST) with specified isotype, numbering scheme, and VJ recombinations, identifying rare SHMs for an input sequence, and showing the GSSP of a V gene (Fig. 5). Users can also query GSNP of an input gene as well as download all the datasets in this study.

**Figure 5.**
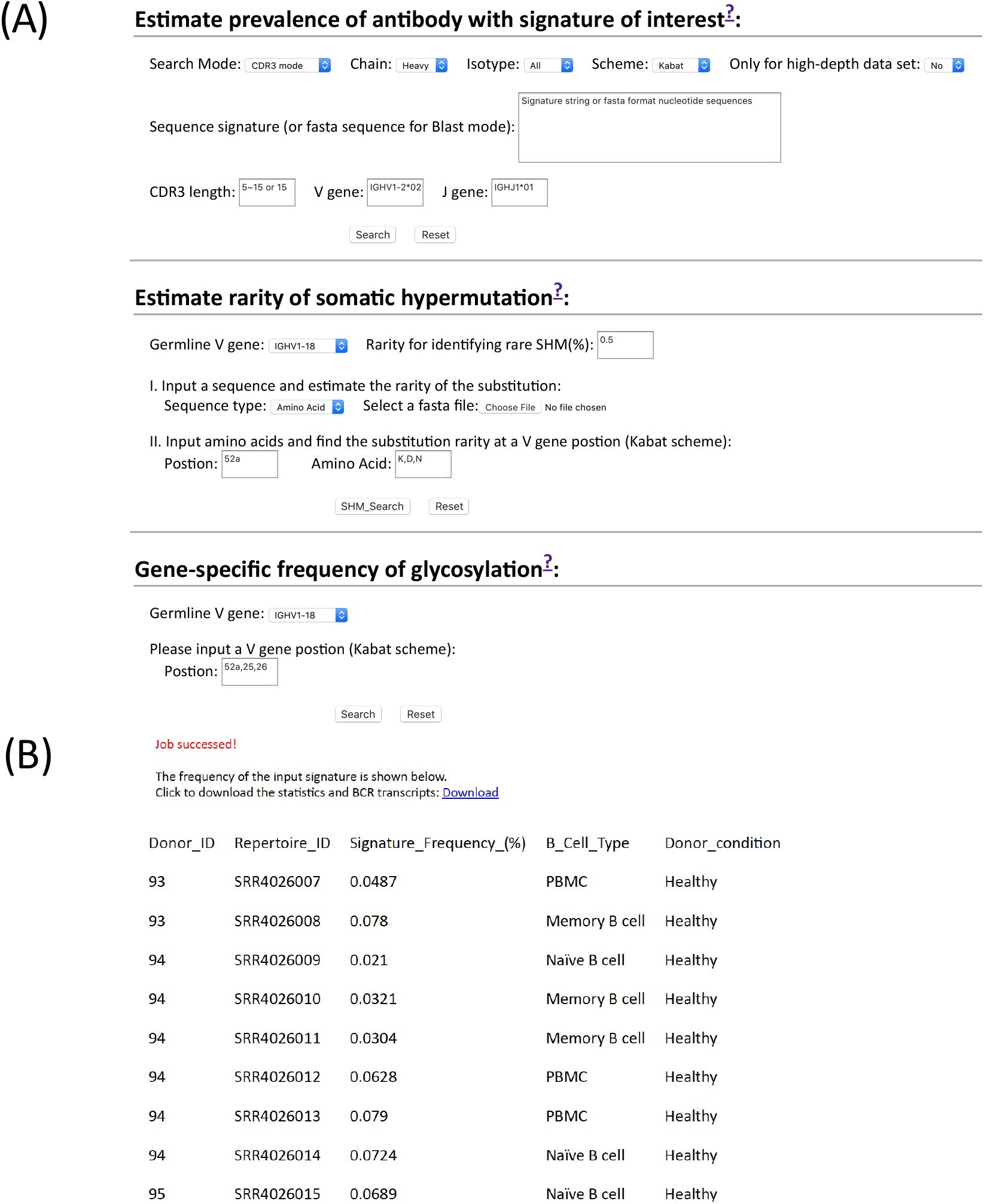
Querying frequencies of signature and somatic hypermutation at cAb-Rep website. (A) cAb-Rep interfaces to estimate frequency of an input signature, gene-specific rarity of somatic hypermutation, and gene-specific glycosylation frequency. (B) Result for VRC01 bnAb-like light chain sequences using the CDR3 mode. The full results including sequences and donor information can be obtained from the download link, and the frequency of the motif in each donor is shown online. The parameters used are: IGKV3-20 gene and a [A-Z]{2}[AFILMYWV][EQ][A-Z] motif.

### Discussion

To understand the mechanisms of BCR diversity generated by V(D)J recombination and somatic hypermutation, high quality BCR repertoire datasets are critical. In this study, we have constructed a database of curated BCR transcripts from previous deep sequencing studies of B cell immunoglobulin sequence repertoires, and demonstrated how this database can provide helpful information for repertoire studies both locally and online. Currently, heavy-light pairing information is not obtained in cAb-Rep datasets. As new technologies are applied to repertoire sequencing, we believe more paired heavy-light chain transcripts will be incorporated, which will greatly advance functional characterization of BCRs. We will continue to update this database to include more disease conditions as well as those from animal models.

In cAb-Rep, we incorporated BCR datasets sequenced both with and without UMI. Besides filtering out transcripts with low sequencing quality, different approaches were used to identify high quality transcripts. For repertoires sequenced using UMI, we used the UMI information to generate consensus sequences, which has been proven effective at removing sequencing errors (Khan et al., 2016). For BCR repertoires sequenced without UMI, we used a clustering approach, which we first sorted transcripts based on redundancy and selected transcripts with the most redundancy as seeds to cluster transcripts at a given identity cutoff and only used the seeds as high quality transcripts. The assumption is that each B cell contains multiple BCR mRNA molecules and the original BCR could be PCR amplified and sequenced with many copies. Sequencing errors and PCR crossover are rare and close to random events (Schirmer et al., 2015), sequences containing sequencing errors are likely a few hamming distance away from the seed transcripts or appear as singletons after clustering. Thus, removing transcripts highly similar to the seed transcripts and singletons will remove many sequencing errors, although we may lose a portion of biological transcripts with low sequencing coverage. This approach is proven effective at finding functional transcripts in previous studies (Bhiman et al., 2015; Krebs et al., 2019).

Compared to other BCR repertoire databases, cAb-Rep provides more flexibility to genetic search signatures. Another public database, iReceptor, supported by the adaptive immune receptor repertoire community, has been developed to store, query, and analyze immune receptor repertoire data (Corrie et al., 2018). While iReceptor includes BCR repertoires amplified with various library preparation protocols and sequenced in multiple platforms (illumine, 454, ion torrent, etc.), cAb-Rep is limited to include full-length or near-full length sequences with enough sample size for repertoire related analyses. For repertoire analysis, iReceptor allows query with V(D)J recombination and a CDR3 peptide (no regular expression grammars allowed), while cAb-Rep provides BLAST mode and a more flexible CDR3 searching mode. cAb-Rep also allows to search motifs across the V(D)J region in the position mode. Although iReceptor includes curated repertoire data deposited by researchers, these curated datasets are processed using different bioinformatics pipeline and parameters (germline gene database, exclusion of redundancy, etc.), which may introduce bias when performing statistical analysis across studies. Another B cell repertoire database is available, but only includes repertoires from three donors (DeWitt et al., 2016). Moreover, the 13 HD curated BCR repertoires, which provide high depth BCR diversity information, haven’t been incorporated in other databases. These datasets increase the statistical power to study features as well as rare events of BCR diversity.

The second goal of cAb-Rep is to advance the investigation of functions of somatic hypermutation, which, as far as we know, is not available in other B cell repertoire databases. We provide a query portal and GSSPs for human antibody V genes to understand gene- and positional-specific substitution preferences. We also predicted gene-specific frequencies of N-glycosylation in human antibody V genes. Such sequence-derived information, together with functional study, is critical to identify common mechanism of SHM functioning and understand similarities in pathways of antibody affinity maturation (Bonsignori et al., 2016; Koenig et al., 2017; van de Bovenkamp et al., 2018). Nonetheless, our current knowledge on how SHM function is still very limited, and a database to comprehensively annotate functions of SHM will accelerate SHM studies. In the future, we will collect literatures to annotate functions of SHM as well as develop new tools to predict functions of SHM. A portal in cAb-Rep will be created to query functions of SHM, which will elucidate the relations of antibody sequence, structure, and function and provide knowledge for antibody design.

In summary, we believe cAb-Rep and the tools developed in this study are complimentary to other B cell repertoire databases, and will be helpful to researchers not familiar with repertoire annotation to explore features of repertoires and compare datasets across studies and diseases.

## Author Contributions

ZS, PK, and LS designed the research; ZS, YG, and KC analyzed data; ZS, YG, LS, and PK wrote the paper. All authors reviewed, commented on, and approved the manuscript.

## References

Altschul, S.F., Madden, T.L., Schaffer, A.A., Zhang, J., Zhang, Z., Miller, W., and Lipman, D.J. (1997). Gapped BLAST and PSI-BLAST: a new generation of protein database search programs. Nucleic Acids Res 25, 3389–3402.

Andrabi, R., Voss, J.E., Liang, C.H., Briney, B., McCoy, L.E., Wu, C.Y., Wong, C.H., Poignard, P., and Burton, D.R. (2015). Identification of Common Features in Prototype Broadly Neutralizing Antibodies to HIV Envelope V2 Apex to Facilitate Vaccine Design. Immunity 43, 959–973.

Bhiman, J.N., Anthony, C., Doria-Rose, N.A., Karimanzira, O., Schramm, C.A., Khoza, T., Kitchin, D., Botha, G., Gorman, J., Garrett, N.J., et al. (2015). Viral variants that initiate and drive maturation of V1V2-directed HIV-1 broadly neutralizing antibodies. Nature medicine 21, 1332–1336.

Bonsignori, M., Zhou, T., Sheng, Z., Chen, L., Gao, F., Joyce, M.G., Ozorowski, G., Chuang, G.Y., Schramm, C.A., Wiehe, K., et al. (2016). Maturation Pathway from Germline to Broad HIV-1 Neutralizer of a CD4-Mimic Antibody. Cell 165, 449–463.

Briney, B., Inderbitzin, A., Joyce, C., and Burton, D.R. (2019). Commonality despite exceptional diversity in the baseline human antibody repertoire. Nature.

Brochet, X., Lefranc, M.P., and Giudicelli, V. (2008). IMGT/V-QUEST: the highly customized and integrated system for IG and TR standardized V-J and V-D-J sequence analysis. Nucleic Acids Res 36, W503–508.

Burton, D.R., and Mascola, J.R. (2015). Antibody responses to envelope glycoproteins in HIV-1 infection. Nature immunology 16, 571–576.

Camacho, C., Coulouris, G., Avagyan, V., Ma, N., Papadopoulos, J., Bealer, K., and Madden, T.L. (2009). BLAST+: architecture and applications. BMC Bioinformatics 10, 421.

Chaudhary, N., and Wesemann, D.R. (2018). Analyzing Immunoglobulin Repertoires. Frontiers in immunology 9, 462.

Chazenbalk, G.D., Portolano, S., Russo, D., Hutchison, J.S., Rapoport, B., and McLachlan, S. (1993). Human organ-specific autoimmune disease. Molecular cloning and expression of an autoantibody gene repertoire for a major autoantigen reveals an antigenic immunodominant region and restricted immunoglobulin gene usage in the target organ. The Journal of clinical investigation 92, 62–74.

Chuang, G.Y., Zhou, J., Acharya, P., Rawi, R., Shen, C.H., Sheng, Z.Z., Zhang, B.S., Zhou, T.Q., Bailer, R.T., Dandey, V.P., et al. (2019). Structural Survey of Broadly Neutralizing Antibodies Targeting the HIV-1 Env Trimer Delineates Epitope Categories and Characteristics of Recognition. Structure 27, 196-+.

Corcoran, M.M., Phad, G.E., Vazquez Bernat, N., Stahl-Hennig, C., Sumida, N., Persson, M.A., Martin, M., and Karlsson Hedestam, G.B. (2016). Production of individualized V gene databases reveals high levels of immunoglobulin genetic diversity. Nature communications 7, 13642.

Corrie, B.D., Marthandan, N., Zimonja, B., Jaglale, J., Zhou, Y., Barr, E., Knoetze, N., Breden, F.M.W., Christley, S., Scott, J.K., et al. (2018). iReceptor: A platform for querying and analyzing antibody/B-cell and T-cell receptor repertoire data across federated repositories. Immunological reviews 284, 24–41.

Corti, D., and Lanzavecchia, A. (2013). Broadly neutralizing antiviral antibodies. Annual review of immunology 31, 705–742.

DeWitt, W.S., Lindau, P., Snyder, T.M., Sherwood, A.M., Vignali, M., Carlson, C.S., Greenberg, P.D., Duerkopp, N., Emerson, R.O., and Robins, H.S. (2016). A Public Database of Memory and Naive B-Cell Receptor Sequences. PLoS One 11, e0160853.

Doria-Rose, N.A., Schramm, C.A., Gorman, J., Moore, P.L., Bhiman, J.N., DeKosky, B.J., Ernandes, M.J., Georgiev, I.S., Kim, H.J., Pancera, M., et al. (2014). Developmental pathway for potent V1V2-directed HIV-neutralizing antibodies. Nature 509, 55–62.

Dosenovic, P., Kara, E.E., Pettersson, A.K., McGuire, A.T., Gray, M., Hartweger, H., Thientosapol, E. S., Stamatatos, L., and Nussenzweig, M.C. (2018). Anti-HIV-1 B cell responses are dependent on B cell precursor frequency and antigen-binding affinity. Proc Natl Acad Sci U S A 115, 4743–4748.

Dunbar, J., and Deane, C.M. (2016). ANARCI: antigen receptor numbering and receptor classification. Bioinformatics 32, 298–300.

Edgar, R.C., and Flyvbjerg, H. (2015). Error filtering, pair assembly and error correction for next-generation sequencing reads. Bioinformatics 31, 3476–3482.

Ekiert, D.C., Bhabha, G., Elsliger, M.A., Friesen, R.H., Jongeneelen, M., Throsby, M., Goudsmit, J., and Wilson, I.A. (2009). Antibody recognition of a highly conserved influenza virus epitope. Science 324, 246–251.

Elhanati, Y., Sethna, Z., Marcou, Q., Callan, C.G., Jr., Mora, T., and Walczak, A.M. (2015). Inferring processes underlying B-cell repertoire diversity. Philos Trans R Soc Lond B Biol Sci 370.

Francica, J.R., Sheng, Z., Zhang, Z., Nishimura, Y., Shingai, M., Ramesh, A., Keele, B.F., Schmidt, S.D., Flynn, B.J., Darko, S., et al. (2015). Analysis of immunoglobulin transcripts and hypermutation following SHIV(AD8) infection and protein-plus-adjuvant immunization. Nature communications 6, 6565.

Galson, J.D., Truck, J., Clutterbuck, E.A., Fowler, A., Cerundolo, V., Pollard, A.J., Lunter, G., and Kelly, D.F. (2016a). B-cell repertoire dynamics after sequential hepatitis B vaccination and evidence for cross-reactive B-cell activation. Genome Med 8, 68.

Galson, J.D., Truck, J., Fowler, A., Clutterbuck, E.A., Munz, M., Cerundolo, V., Reinhard, C., van der Most, R., Pollard, A.J., Lunter, G., and Kelly, D.F. (2015). Analysis of B Cell Repertoire Dynamics Following Hepatitis B Vaccination in Humans, and Enrichment of Vaccine-specific Antibody Sequences. EBioMedicine 2, 2070–2079.

Galson, J.D., Truck, J., Kelly, D.F., and van der Most, R. (2016b). Investigating the effect of AS03 adjuvant on the plasma cell repertoire following pH1N1 influenza vaccination. Sci Rep 6, 37229.

Gupta, N.T., Adams, K.D., Briggs, A.W., Timberlake, S.C., Vigneault, F., and Kleinstein, S.H. (2017). Hierarchical Clustering Can Identify B Cell Clones with High Confidence in Ig Repertoire Sequencing Data. Journal of Immunology 198, 2489–2499.

Hexham, J.M., Furmaniak, J., Pegg, C., Burton, D.R., and Smith, B.R. (1992). Cloning of a human autoimmune response: preparation and sequencing of a human anti-thyroglobulin autoantibody using a combinatorial approach. Autoimmunity 12, 135–141.

Imkeller, K., Scally, S.W., Bosch, A., Marti, G.P., Costa, G., Triller, G., Murugan, R., Renna, V., Jumaa, H., Kremsner, P.G., et al. (2018). Antihomotypic affinity maturation improves human B cell responses against a repetitive epitope. Science 360, 1358–1362.

Jardine, J.G., Kulp, D.W., Havenar-Daughton, C., Sarkar, A., Briney, B., Sok, D., Sesterhenn, F., Ereno-Orbea, J., Kalyuzhniy, O., Deresa, I., et al. (2016a). HIV-1 broadly neutralizing antibody precursor B cells revealed by germline-targeting immunogen. Science 351, 1458–1463.

Jardine, J.G., Sok, D., Julien, J.P., Briney, B., Sarkar, A., Liang, C.H., Scherer, E.A., Henry Dunand, C.J., Adachi, Y., Diwanji, D., et al. (2016b). Minimally Mutated HIV-1 Broadly Neutralizing Antibodies to Guide Reductionist Vaccine Design. PLoS pathogens 12, e1005815.

Jayaram, N., Bhowmick, P., and Martin, A.C. (2012). Germline VH/VL pairing in antibodies. Protein Eng Des Sel 25, 523–529.

Jeon, Y.E., Seo, C.W., Yu, E.S., Lee, C.J., Park, S.G., and Jang, Y.J. (2007). Characterization of human monoclonal autoantibody Fab fragments against oxidized LDL. Mol Immunol 44, 827–836.

Joyce, M.G., Wheatley, A.K., Thomas, P.V., Chuang, G.Y., Soto, C., Bailer, R.T., Druz, A., Georgiev, I.S., Gillespie, R.A., Kanekiyo, M., et al. (2016). Vaccine-Induced Antibodies that Neutralize Group 1 and Group 2 Influenza A Viruses. Cell 166, 609–623.

Khan, T.A., Friedensohn, S., de Vries, A.R., Straszewski, J., Ruscheweyh, H.J., and Reddy, S.T. (2016). Accurate and predictive antibody repertoire profiling by molecular amplification fingerprinting. Sci Adv 2, e1501371.

Koenig, P., Lee, C.V., Walters, B.T., Janakiraman, V., Stinson, J., Patapoff, T.W., and Fuh, G. (2017). Mutational landscape of antibody variable domains reveals a switch modulating the interdomain conformational dynamics and antigen binding. Proc Natl Acad Sci U S A 114, E486–E495.

Krebs, S.J., Kwon, Y.D., Schramm, C.A., Law, W.H., Donofrio, G., Zhou, K.H., Gift, S., Dussupt, V., Georgiev, I.S., Schätzle, S., et al. (2019). Longitudinal Analysis Reveals Early Development of Three MPER-Directed Neutralizing Antibody Lineages from an HIV-1-Infected Individual. Immunity 50, 677–691.e613.

Kwong, P.D., Chuang, G.Y., DeKosky, B.J., Gindin, T., Georgiev, I.S., Lemmin, T., Schramm, C.A., Sheng, Z., Soto, C., Yang, A.S., et al. (2017). Antibodyomics: bioinformatics technologies for understanding B-cell immunity to HIV-1. Immunological reviews 275, 108–128.

Kwong, P.D., and Mascola, J.R. (2018). HIV-1 Vaccines Based on Antibody Identification, B Cell Ontogeny, and Epitope Structure. Immunity 48, 855–871.

Kwong, P.D., and Wilson, I.A. (2009). HIV-1 and influenza antibodies: seeing antigens in new ways. Nature immunology 10, 573–578.

Margreitter, C., Lu, H.C., Townsend, C., Stewart, A., Dunn-Walters, D.K., and Fraternali, F. (2018). BRepertoire: a user-friendly web server for analysing antibody repertoire data. Nucleic Acids Res 46, W264–W270.

Miho, E., Yermanos, A., Weber, C.R., Berger, C.T., Reddy, S.T., and Greiff, V. (2018). Computational Strategies for Dissecting the High-Dimensional Complexity of Adaptive Immune Repertoires. Frontiers in immunology 9, 224.

Murphy, K. (2014). Janeway’s Immunobiology (Garland Science).

Nielsen, S.C.A., and Boyd, S.D. (2018). Human adaptive immune receptor repertoire analysis-Past, present, and future. Immunological reviews 284, 9–23.

Odegard, V.H., and Schatz, D.G. (2006). Targeting of somatic hypermutation. Nature reviews. Immunology 6, 573–583.

Pappas, L., Foglierini, M., Piccoli, L., Kallewaard, N.L., Turrini, F., Silacci, C., Fernandez-Rodriguez, B., Agatic, G., Giacchetto-Sasselli, I., Pellicciotta, G., et al. (2014). Rapid development of broadly influenza neutralizing antibodies through redundant mutations. Nature 516, 418-+.

Pichurin, P., Guo, J., Yan, X., Rapoport, B., and McLachlan, S.M. (2001). Human monoclonal autoantibodies to B-cell epitopes outside the thyroid peroxidase autoantibody immunodominant region. Thyroid 11, 301–313.

Pichurin, P.N., Guo, J., Estienne, V., Carayon, P., Ruf, J., Rapoport, B., and McLachlan, S.M. (2002). Evidence that the complement control protein-epidermal growth factor-like domain of thyroid peroxidase lies on the fringe of the immunodominant region recognized by autoantibodies. Thyroid 12, 1085–1095.

R. Gupta, E. Jung, and Brunak, S. (2004). Prediction of N-glycosylation sites in human proteins. In Preparation.

Rubelt, F., Bolen, C.R., McGuire, H.M., Vander Heiden, J.A., Gadala-Maria, D., Levin, M., Euskirchen, G.M., Mamedov, M.R., Swan, G.E., Dekker, C.L., et al. (2016). Individual heritable differences result in unique cell lymphocyte receptor repertoires of naive and antigen-experienced cells. Nature communications 7, 11112.

Schirmer, M., Ijaz, U.Z., D’Amore, R., Hall, N., Sloan, W.T., and Quince, C. (2015). Insight into biases and sequencing errors for amplicon sequencing with the Illumina MiSeq platform. Nucleic Acids Research 43, e37.

Schramm, C.A., Sheng, Z., Zhang, Z., Mascola, J.R., Kwong, P.D., and Shapiro, L. (2016). SONAR: A High-Throughput Pipeline for Inferring Antibody Ontogenies from Longitudinal Sequencing of B Cell Transcripts. Frontiers in immunology 7, 372.

Sheng, Z., Schramm, C.A., Connors, M., Morris, L., Mascola, J.R., Kwong, P.D., and Shapiro, L. (2016). Effects of Darwinian Selection and Mutability on Rate of Broadly Neutralizing Antibody Evolution during HIV-1 Infection. PLoS Comput Biol 12, e1004940.

Sheng, Z., Schramm, C.A., Kong, R., Program, N.C.S., Mullikin, J.C., Mascola, J.R., Kwong, P.D., Shapiro, L., Benjamin, B., Bouffard, G., et al. (2017). Gene-Specific Substitution Profiles Describe the Types and Frequencies of Amino Acid Changes during Antibody Somatic Hypermutation. Frontiers in immunology 8, 537.

Sievers, F., Wilm, A., Dineen, D., Gibson, T.J., Karplus, K., Li, W., Lopez, R., McWilliam, H., Remmert, M., Soding, J., et al. (2011). Fast, scalable generation of high-quality protein multiple sequence alignments using Clustal Omega. Molecular systems biology 7, 539.

Soto, C., Bombardi, R.G., Branchizio, A., Kose, N., Matta, P., Sevy, A.M., Sinkovits, R.S., Gilchuk, P., Finn, J.A., and Crowe, J.E., Jr. (2019). High frequency of shared clonotypes in human B cell receptor repertoires. Nature 566, 398–402.

Stern, J.N., Yaari, G., Vander Heiden, J.A., Church, G., Donahue, W.F., Hintzen, R.Q., Huttner, A.J., Laman, J.D., Nagra, R.M., Nylander, A., et al. (2014). B cells populating the multiple sclerosis brain mature in the draining cervical lymph nodes. Science translational medicine 6, 248ra107.

Tipton, C.M., Fucile, C.F., Darce, J., Chida, A., Ichikawa, T., Gregoretti, I., Schieferl, S., Hom, J., Jenks, S., Feldman, R.J., et al. (2015). Diversity, cellular origin and autoreactivity of antibody-secreting cell population expansions in acute systemic lupus erythematosus. Nature immunology 16, 755–765.

van de Bovenkamp, F.S., Derksen, N.I.L., Ooijevaar-de Heer, P., van Schie, K.A., Kruithof, S., Berkowska, M.A., van der Schoot, C.E., IJspeert, H., van der Burg, M., Gils, A., et al. (2018). Adaptive antibody diversification through N-linked glycosylation of the immunoglobulin variable region. P Natl Acad Sci USA 115, 1901–1906.

Vander Heiden, J.A., Marquez, S., Marthandan, N., Bukhari, S.A.C., Busse, C.E., Corrie, B., Hershberg, U., Kleinstein, S.H., Matsen Iv, F.A., Ralph, D.K., et al. (2018). AIRR Community Standardized Representations for Annotated Immune Repertoires. Frontiers in immunology 9, 2206.

Vander Heiden, J.A., Stathopoulos, P., Zhou, J.Q., Chen, L., Gilbert, T.J., Bolen, C.R., Barohn, R.J., Dimachkie, M.M., Ciafaloni, E., Broering, T.J., et al. (2017). Dysregulation of B Cell Repertoire Formation in Myasthenia Gravis Patients Revealed through Deep Sequencing. Journal of Immunology 198, 1460–1473.

West, A.P., Jr., Diskin, R., Nussenzweig, M.C., and Bjorkman, P.J. (2012). Structural basis for germ-line gene usage of a potent class of antibodies targeting the CD4-binding site of HIV-1 gp120. Proc Natl Acad Sci U S A 109, E2083–2090.

Wu, X., Liu, B., Van der Merwe, P.L., Kalis, N.N., Berney, S.M., and Young, D.C. (1998). Myosin-reactive autoantibodies in rheumatic carditis and normal fetus. Clinical immunology and immunopathology 87, 184–192.

Wu, X., Zhang, Z., Schramm, C.A., Joyce, M.G., Kwon, Y.D., Zhou, T., Sheng, Z., Zhang, B., O’Dell, S., McKee, K., et al. (2015). Maturation and Diversity of the VRC01-Antibody Lineage over 15 Years of Chronic HIV-1 Infection. Cell 161, 470–485.

Wu, X., Zhou, T., Zhu, J., Zhang, B., Georgiev, I., Wang, C., Chen, X., Longo, N.S., Louder, M., McKee, K., et al. (2011). Focused evolution of HIV-1 neutralizing antibodies revealed by structures and deep sequencing. Science 333, 1593–1602.

Zhou, T., Lynch, R.M., Chen, L., Acharya, P., Wu, X., Doria-Rose, N.A., Joyce, M.G., Lingwood, D., Soto, C., Bailer, R.T., et al. (2015). Structural Repertoire of HIV-1-Neutralizing Antibodies Targeting the CD4 Supersite in 14 Donors. Cell 161, 1280–1292.

Zhou, T., Zhu, J., Wu, X., Moquin, S., Zhang, B., Acharya, P., Georgiev, I.S., Altae-Tran, H.R., Chuang, G.Y., Joyce, M.G., et al. (2013). Multidonor analysis reveals structural elements, genetic determinants, and maturation pathway for HIV-1 neutralization by VRC01-class antibodies. Immunity 39, 245–258.

